# Dynamic structural connectivity changes in cortical and cortico-striatal strokes in mice

**DOI:** 10.1101/2025.03.27.644898

**Authors:** Fatemeh Mahani, Aref Kalantari, Michael Diedenhofen, Claudia Green, Dirk Wiedermann, Gereon R. Fink, Mathias Hoehn, Markus Aswendt

## Abstract

Stroke is a primary global health concern, leading to significant mortality and long-term disability. Beyond immediate neuronal damage, functional and structural connectivity is altered brain-wide with implications for functional deficits and recovery. It remains unclear, however, if the level of degeneration, i.e. reduced myelination and axonal damage, as well as compensatory plasticity, i.e., axonal sprouting and remyelination, depend on the lesion size and topology. This study compares for the first time two different stroke models in adult male mice, with the aim of uncovering the dynamics in white matter changes. Repetitive diffusion magnetic resonance imaging (dMRI) over four weeks post photothrombotic cortical (1.41±0.92% of brain volume), and middle cerebral artery occlusion (MCAO) cortico-striatal (11.53±2.8% of brain volume), strokes were used to map structural connectivity changes at the whole-brain level. We quantified inter- and intra-hemispheric seed strength changes over time, with seed strength reflecting how strongly each region is connected to the rest of the brain. Differences between groups and time points were assessed using a mixed model corrected for multiple comparisons. In conclusion, large cortico-striatal lesions led to increased structural connectivity in sensorimotor regions, whereas small cortical lesions induced asymmetric connectivity changes: an increase extending globally from the ischemic hemisphere and a decrease expanding globally from the healthy hemisphere. These findings highlight that stroke severity and lesion size significantly affect the temporal dynamics and spatial distribution of connectivity disruptions, emphasizing the need for targeted monitoring of neural changes post-stroke.

**Significance statement:** Pathophysiological changes close to an ischemic lesion are well understood. But structurally connected regions also undergo stroke-related changes potentially contributing to functional deficits. Animal studies using in vivo diffusion MRI to quantify white matter changes have been limited to single connections, time points after stroke, and basic diffusion measures. We provide the first in-depth characterization of whole-brain fiber tracking to compare small cortical and large cortico-striatal strokes. We found a complex dynamic of decreasing and increasing structural connectivity in single sensorimotor-related brain regions within the brain network - even remote areas from the stroke lesion. This study provides a better understanding of the interactions of regions during stroke recovery, which may help in developing therapies targeting localized white matter repair.

## Introduction

A stroke results in significant impairment of motor and cognitive functions and is a leading cause of long-term disability worldwide. Early after stroke, the brain undergoes a cascade of pathophysiological events that significantly alter its structural and functional organization in the ischemic territory and beyond (Dirnagl et al., 1999; Carmichael, 2016). In the days and weeks post-stroke, secondary processes such as gliosis, axonal degeneration, and synaptic remodeling take place, contributing to both local and widespread network alterations (Grefkes and Fink, 2014; Pekny and Pekna, 2014; Carmichael et al., 2017). These pathological changes disrupt connections disintegrating functional networks, but simultaneously stimulate compensatory mechanisms aimed at restoring connectivity through synaptic plasticity, axonal sprouting, and the recruitment of alternate pathways (Pekny and Pekna, 2014). This dynamic reorganization is crucial for functional recovery, as it enables the remaining healthy networks to adapt and compensate for the lost connections, ultimately shaping the extent of rehabilitation (Jones, 2017). Understanding how these networks change over time is critical for developing effective rehabilitation strategies (Grefkes and Fink, 2014). Diffusion MRI (dMRI) is to date the most widely used method to quantify white matter changes in relation to motor deficit and recovery after stroke longitudinally (Moura et al., 2019). Preclinical studies in rodents are lacking to fully capture the mechanistic principles of white matter changes and their relevance for motor recovery. Earlier studies were limited to short time windows of acute stroke (Cha et al., 2016; Chang et al., 2017; Chiu et al., 2018; Alves et al., 2022) or individual time points after stroke induction (Li et al., 2020; Alves et al., 2022), or have not included pre-stroke control data as would be helpful in direct reference with non-invasive imaging approaches (Li et al., 2020). Often, only selective and specific connections of white matter structures were analyzed. Here, typically, a primary focus was limited to aspects of large white matter bundles of the internal and external capsules, and corpus callosum (van Meer et al., 2011; Po et al., 2012; Zanier et al., 2013; Cha et al., 2016; Jung et al., 2017; Sinke et al., 2018; Anon, 2020). Thus, previous rodent dMRI studies provided detailed information about individual connections but did not comprehensively inform about the wide-spread structural changes across hemispheres. This study investigated the time-dependent changes in fiber density within and between brain regions during the recovery phase after permanent cortical stroke and transient cortico-striatal stroke in mice. Specifically, we focused on the sensorimotor, thalamic, and white matter networks, which are known to play key roles in functional recovery post-stroke. Using longitudinal scans over four weeks post-stroke, fiber density was calculated, together with seed strength, a measure of fiber density that reflects the connectivity of a brain region relative to the rest of the brain. This approach allowed the mapping of intra- and inter-hemispheric structural connections over time, providing insight into the dynamics of network reorganization across different phases of recovery in two different stroke models.

## Materials and Methods

### Experimental design

Longitudinal in vivo MRI data were derived from pre-existing studies, which were part of previous publications (Green et al., 2018; Pallast et al., 2020; Aswendt et al., 2021). These datasets represent two distinct mouse models of stroke: a cortical lesion model generated using the photothrombosis technique and a cortico-striatal lesion model produced by the MCAO (middle cerebral artery occlusion) method in adult mice. All animal procedures were conducted under the German Animal Welfare Act and approved by the Landesamt für Natur, Umwelt und Verbraucherschutz North Rhine-Westphalia. Mice were housed under a 12-hour light/dark cycle with access to food and water. Both datasets, previously collected and repurposed for the current analysis, consist of structural magnetic resonance imaging (MRI) data, with experiments adhering to the ARRIVE and IMPROVE guidelines. Imaging was performed using a 94/20USR BioSpec Bruker system (Bruker BioSpin, Ettlingen, Germany) equipped with a cryo-coil and operated under ParaVision (v6.0.1). Mice were anesthetized with isoflurane (2-3% in 70/30 N2/O2) and stabilized in an animal carrier. Vital parameters, including respiration and body temperature, were monitored throughout the procedure to ensure animal welfare and stable physiological condition throughout the MR session. A T2-weighted RARE sequence was acquired to obtain high-resolution, whole-brain images, followed by diffusion imaging to assess structural connectivity changes.

#### Cortical stroke dataset

Photothrombosis to induce cortical stroke was applied in N=25 male C57Bl/6J mice (8-10 weeks old), following the protocol described in (Aswendt et al., 2021). Briefly, after the injection of Rose Bengal, a photosensitive dye, targeted laser irradiation to the cortex caused localized endothelial damage and thrombus formation. Anatomical T2-weighted MRI and structural DTI (**Table 1**) were performed before stroke (baseline) and at week 1, 2, and 4 post-stroke.

**Table 1:** MRI parameters.

| Study | Sequence | TR (ms) | TE (ms) | Acq. time (min) | Voxel ($\mu\text{m}^3$) | FOV ($\text{mm}^2$) | Flip angle | B-value ($\text{s}/\text{mm}^2$) | Diffusion directions |
| --- | --- | --- | --- | --- | --- | --- | --- | --- | --- |
| PT | Turbo-RARE | 5.5 | 32.5 | 5:52 | 68.4×68.4×300 | 17.5×17.5 | 90 | ~ | ~ |
|  | DTI | 3.0 | 17.5 | 14:00 | 141×141×400 | 18×18 | 90 | 677 | 30 |
| MCAO | Turbo-RARE | 5.5 | 32.5 | 4:58 | 68.4×68.4×200 | 17.5×17.5 | 180 | ~ | ~ |
|  | DSI | 3.5 | 20s | 3:58 | 139×139×500 | 17.8×17.8 | 90 | 2,000 | 126<sup>1</sup> |

<sup>1</sup> half square

#### Cortico-striatal stroke dataset

This dataset involved a transient middle cerebral artery occlusion (MCAO) stroke model in N=6 male NMRI-nu mice (12–14 weeks old, 32–37 g), as described in (Green et al., 2018). Briefly, MCAO was induced by inserting a rubber-coated filament into the internal carotid artery for 30 minutes, followed by reperfusion. Anatomical T2-weighted and structural diffusion-sensitive Q-ball MR imaging (**Table 1**) were acquired before stroke (baseline) and at week 1, 2, and 4 post-stroke.

### MRI processing

MRI data were processed using **AIDAmri** (version 1.3) pipeline (Pallast et al., 2019) and registered with the Allen Mouse Brain Reference Atlas (AMBA, CCF v3.0).

#### Pre-processing and atlas registration

Pre-processing of the data included conversion from Bruker raw data format to NIfTI format, brain extraction (using FSL bet), and bias field correction (Multiplicative Intrinsic Component Optimization algorithm as described by (Li et al., 2014)). The registration followed a multi-step procedure of affine and non-linear registration of the T2-weighted MRI with the AMBA first, followed by the DTI registration to the T2-weighted data.

#### Lesion segmentation

Ischemic stroke lesions, identifiable as hyperintense regions on the T2-weighted MRI, were semi-automatically segmented using the **ITK-SNAP** software (Yushkevich et al., 2006). The 3D snake evolution tool within ITK-SNAP enabled precise delineation of lesion boundaries. The resulting stroke masks are transformed to the AMBA, allowing for qualitative and quantitative comparisons of ischemic tissue across different groups, utilizing incidence maps and region-specific lesion calculations, respectively.

#### Tractography and connectivity analysis

Whole-brain tractography and structural connectivity analysis were performed using **DSI Studio version 2023** (Yeh et al., 2013) following 4 consecutive steps: **1) Fiber Tracking:** Deterministic fiber tracking was conducted across the entire brain using a modified AMBA template encompassing 98 selected regions of interest (ROIs). The tracking process utilized parameters such as fiber length of >0.5 or <120 mm, turning angle of <55°, and tracking interval to ensure accurate reconstruction of brain fiber tracts. For the DTI-based dataset, 30 gradient directions were used, and for the DSI-based dataset, 126 unique diffusion directions were employed, which were distributed on a half-sphere in q-space. **2) Adjacency Matrix Construction:** The tractography output was an adjacency matrix, where each entry represented the number of fibers connecting two brain regions. This matrix served as the foundation for subsequent graph-theoretical analyses of the brain’s structural connectivity. **3) Calculation of fiber density:** Weights were assigned to connections within the adjacency matrix to calculate fiber density (FD). The FD was computed following the approach proposed by (Hagmann et al., 2008), with modifications to account for region size and fiber length, thereby eliminating biases towards larger regions or longer fibers. **4) Calculation of seed strength:** Seed strength was calculated to quantify the average connectivity of each brain region relative to the rest of the brain in both the ischemic and contralateral hemispheres **(Fig. 1)**. Using the fiber density matrix, the fiber density values for intra-hemispheric and inter-hemispheric connections were extracted and averaged for each region. This metric was computed at multiple time points to evaluate the dynamic changes in structural connectivity over stroke recovery **(Fig. 1)**.

**Figure 1.**
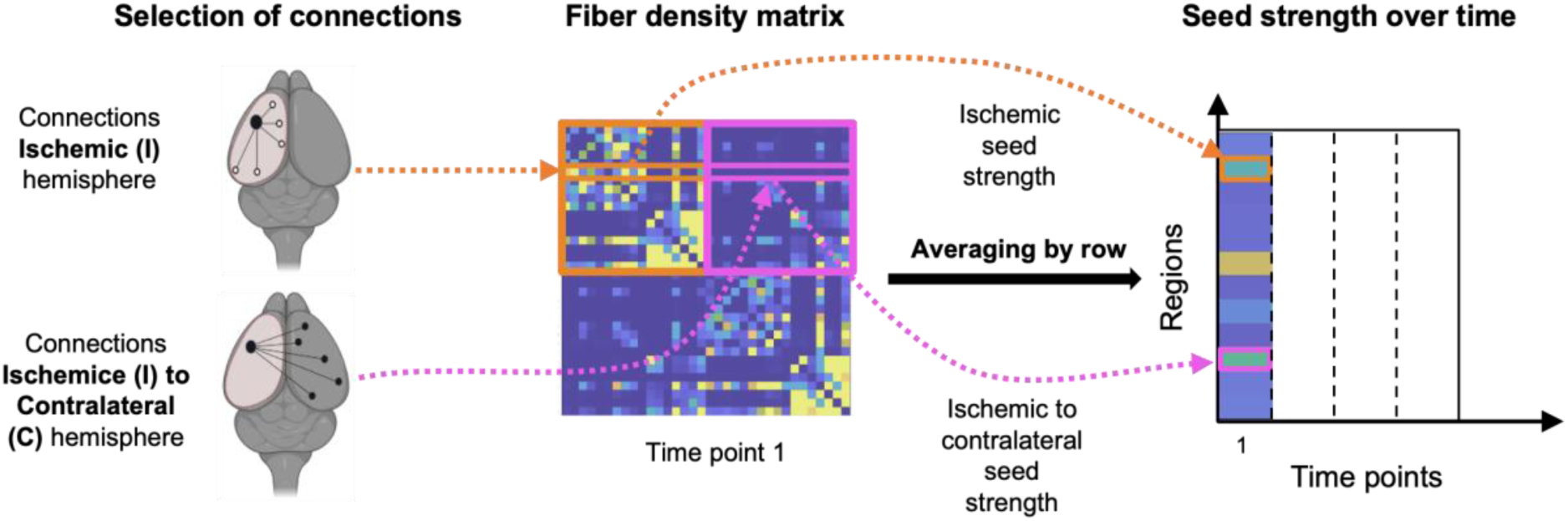
Schematic representation of the calculation of seed strength of individual brain regions to all other brain regions in both ischemic (transparent red) and contralateral (gray) hemispheres. The analysis separated intra-hemispheric and inter-hemispheric connections to assess the average connection strength of each region. Seed strength was calculated using the fiber density matrix of structural brain connections, where the fiber density between each pair of regions is specified. Intra-hemispheric connections (orange rectangle) and inter-hemispheric connections (pink rectangle) are selected for each region, and the fiber density values are averaged across animals to determine the seed strength. The resulting time-series diagram illustrates how the seed strength was built over time based on structural connectivity matrices derived from scans taken at different time points.

#### Excluding the stroke lesion from diffusion MRI processing

Stroke-induced brain edema, caused by fluid accumulation, can significantly distort diffusion imaging and structural network analyses, making it challenging to assess structural connectivity accurately. Before performing fiber tractography, we implemented a method to exclude the stroke lesion masks (initially used to quantify the lesion size and topology) from the fiber orientation distribution (FOD) maps. First, the stroke masks from week 1 were registered to the diffusion imaging scans at all other time points. The transformed lesion areas were set to zero and applied to FOD maps for each time point before proceeding with fiber tracking in DSI Studio. Following this adjustment, we generated updated fiber count matrices and recalculated the volumes of regions partially affected by lesion exclusion. These corrected volume sizes were then used to compute fiber density matrices.

#### Selected brain networks

We analyzed dynamic structural connectivity changes across specific brain networks in mice following cortical and cortico-striatal strokes. The selected brain regions for analysis were based on their known roles in sensorimotor processing and their relevance to stroke-related connectivity changes. The following regions were included in the analysis: **Primary somatosensory cortex (SSp)**: barrel field (**SSp-bfd**), lower limb (**SSp-ll**), mouth (**SSp-m**), nose (**SSp-n**), trunk (**SSp-tr**), upper limb (**SSp-ul**), unidentified area (**SSp-un**), and supplemental somatosensory region (**SSs**); **Motor cortex**: primary motor area (**MOp**) and secondary motor area (**MOs**); **Thalamus (TH)**: postero-medial (**DORpm**), which is associated with the polymodal association cortex, and somatosensory-medial (**DORsm**), which is related to sensory-motor cortical areas; **Striatum:** dorsal striatum (**STRd**) and ventral striatum (**STRv**); **White matter**: corpus callosum (**cc**) and corticospinal tract (**cst**). The SM and TH regions are central to coordinating motor and sensory functions profoundly impacted by stroke. In contrast, the STR regions are integral components of the cortico-striatal circuitry and are highly affected in the cortico-striatal lesion model. The CC and CST are the main inter-hemispheric connection and major motor output. The striatum plays a pivotal role in motor function, reward processing, and neural plasticity, making it a critical focus for understanding connectivity changes in cortico-striatal lesions (Liu et al., 2023). The CC and CST, in particular, mediate compensatory pathways and facilitate neural plasticity, further contributing to recovery dynamics in the mouse brain (Ueno et al., 2012). These regions were selected to comprehensively assess cortical and subcortical areas and white matter pathways to investigate stroke-induced changes in structural connectivity. The connectivity analysis quantified alterations within and between these regions over time, providing insights into the dynamic reorganization of brain networks in response to cortical and cortico-striatal damage.

#### Statistical analyses

A linear mixed-effects model was used to analyze significant changes in seed strength values over time compared to the baseline. The analysis was performed using the lmer function from the lme4 package in R (v4.4.1). The model was specified as follows:

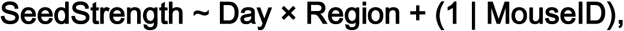

where **FiberDensity** was the dependent variable, **Day** and **Region** were included as fixed effects to capture the influence of time and spatial region, and **MouseID** was treated as a random effect to account for repeated measures within the same mouse. The interaction term (**Day × Region**) allowed for testing whether the effect of time varied by region. The response variable (**SeedStrength**) represents the averaged density of fibers measured in ischemic and contralateral regions over several days, and the inclusion of random intercepts for **MouseID** accommodates inter-subject variability.

Post-hoc pairwise comparisons were performed using **estimated marginal means (EMMs)** via the emmeans package to assess significant changes over time within each region. Pairwise contrasts were evaluated for differences across levels of **Day**, conditional on each level of **Region**. This approach allowed for localized comparisons within each region without collapsing across regions. We applied a correction to account for multiple comparisons and ensure reliable results by controlling the family-wise error rate (FWER). To do this, we used the **Tukey adjustment**, which is particularly effective when comparing all pairs of groups. This method helps maintain accuracy by reducing the chance of false positives while considering how the comparisons are connected.

All statistical analyses were conducted in R (v4.4.1) for macOS using these packages: 1) **lme4**: For fitting the linear mixed-effects model. 2) **emmeans**: For computing estimated marginal means and performing pairwise contrasts. 3) **dplyr**: For data manipulation and pre-processing.

## Results

### Ischemic lesion quantification

The cortical and cortico-striatal lesions, as determined 1 week post-stroke using T2-weighted MRI differed significantly (p<0.001) in overall lesion volume (1.41±0.92% and 11.53±2.8%, respectively) as well as lesion topology (**Fig. 2**). The quantitative lesion mapping revealed expected differences between the stroke models with MCAO leading to a more wide-spread ischemic lesion extending towards the distal somatosensory regions as well as subcortical regions compared to photothrombosis. The atlas-based quantification revealed significant differences in cortical and sub-cortical regions. Cortical lesions covered more area of the primary somatosensory regions (e.g. upper limb, SSp-ul, p<0.01, and lower limb, SSp-ll, p<0.001). In contrast, the cortico-striatal lesions affected to a larger percentage more distal primary and secondary somatosensory regions (e.g. barrel field, SSp-bfd, p<0.001), as well as the dorsal and ventral part of the striatum (STRd and STRv, both p<0.001). No significant difference was detected for the motor cortex and the white matter regions (corticospinal tract, corpus callosum, and medial forebrain bundle system), which were affected to a much lower extent compared to the cortical regions.

**Figure 2.**
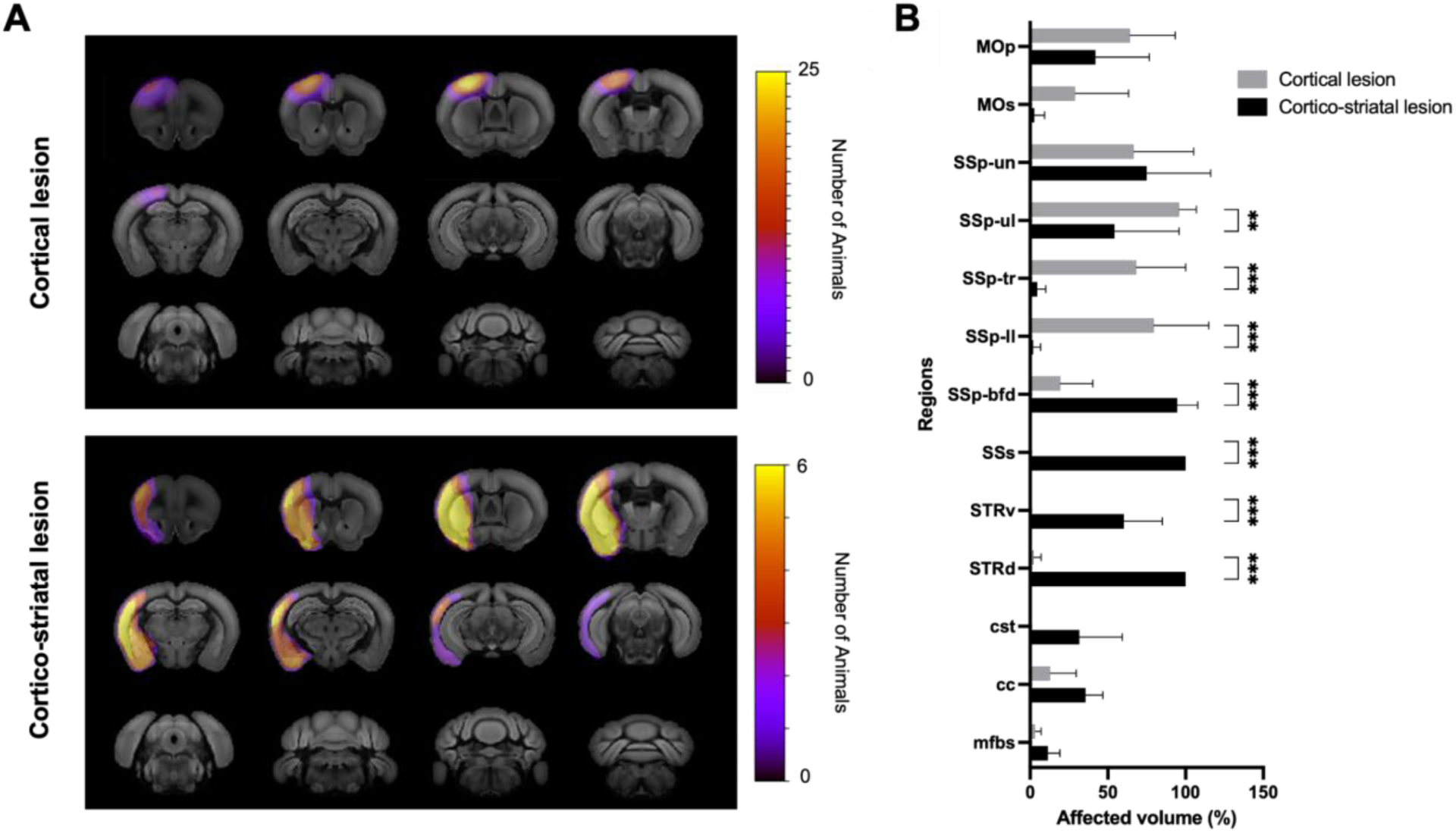
Qualitative and quantitative comparison of lesion size and topology for the cortical and cortico-striatal stroke model. A) Incidence maps quantifying the voxel inside the stroke mask was averaged across mice and transferred back to the atlas reference space to qualitatively compare stroke lesions for both stroke groups. B) Quantitative comparison of affected volume per brain region as determined by individual atlas-based quantification of stroke lesion mask overlap with atlas regions.

### Fiber density

We next asked how these differences in lesion size and topology were reflected in the structural network during the observation time of four weeks post stroke. Individual structural connectivity data were available for 98 atlas regions, i.e., 2401 connections for four time points, i.e., pre-stroke baseline, and 1, 2, and 4 weeks post-stroke. We therefore limited the analysis to the sensorimotor (cortex - striatum - thalamus) system and related white matter regions (corpus callosum and corticospinal tract). Fiber density matrices (as explained in **Fig. 1**) indicating the difference to baseline revealed significant differences between the two stroke models (**Fig. 3**). **Cortical lesion group:** In both hemispheres, approximately 60% of intra-hemispheric connections decreased during the first week after stroke. However, at 2 and 4 weeks, a significant recovery was observed in the ischemic hemisphere, with about 50% of the connections showing an increase. In contrast, in the contralateral hemisphere, the white matter (WM) and striatum (STR) regions demonstrated the largest contribution to the observed increase in connections at 2 and 4 weeks post-stroke. The contralesional sensorimotor and thalamic regions exhibited a decrease in fiber density compared to baseline and the ischemic hemisphere. **Cortico-striatal lesion group:** In the ischemic hemisphere, sensorimotor (SM) and thalamic (TH) regions demonstrated a consistent increase of approximately 25% in intra-hemispheric connections over time. In contrast, the striatum (STR) and white matter (WM) regions of the ischemic hemisphere exhibited a sustained decrease in connectivity. In the contralateral hemisphere, the sensorimotor (SM) and thalamic (TH) regions displayed a stable 40% increase starting at week 1 in intra-hemispheric connections. Similarly, the white matter regions of this hemisphere showed a gradual increase in connectivity over time.

**Figure 3:**
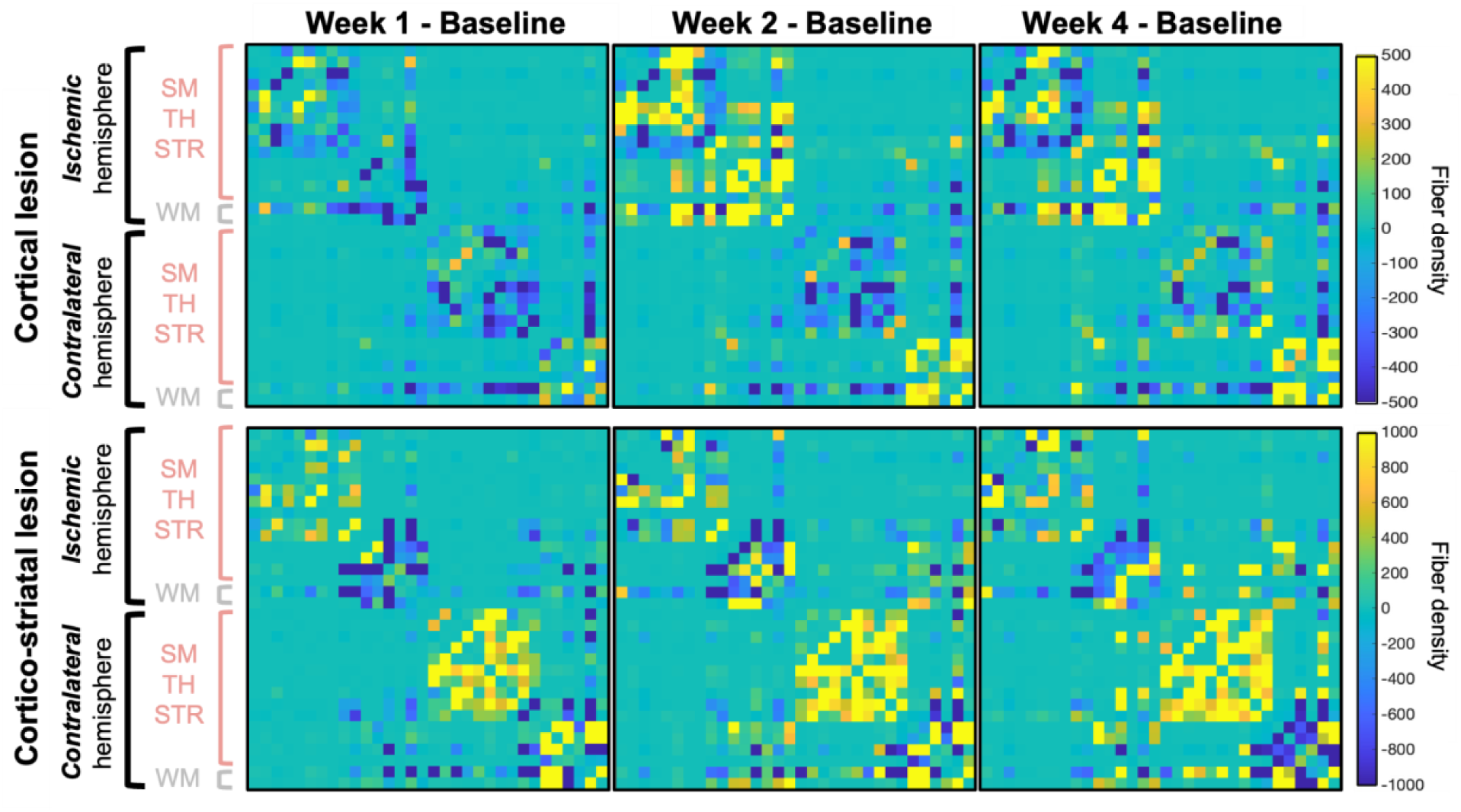
Variability in fiber density changes relative to baseline in the cortical and cortico-striatal stroke model. The matrices represent structural connections in the sensorimotor networks including the motor and sensory cortex (SM), thalamus (TH), and striatum (STR), as well as white matter (WM) tracts at 1, 2, and 4 weeks post-stroke. Fiber density values were normalized relative to the pre-stroke baseline levels. Yellow values in the matrices indicate an increase in fiber density, while blue values represent a decrease.

In summary, the fiber density analysis revealed distinct connectivity patterns between the two stroke models, with cortical lesions showing initial reductions followed by recovery in the ischemic hemisphere and cortico-striatal lesions demonstrating consistent increases in specific regions across both hemispheres.

### Seed strength

Based on these pronounced differences in fiber density between the two stroke models, we were interested in the dynamic changes of individual seeds on the whole brain. Therefore, we calculated the overall connectivity strength of each region, referred to as seed strength, which represents the sum of fiber densities with all other connected regions (**Fig. 4**). Using a linear mixed model with corrections for multiple comparisons, significant temporal changes and regional differences in connectivity within the sensorimotor and thalamic regions were identified. These changes were observed within the ischemic and contralateral hemisphere, respectively, and in inter-hemispheric connections in both stroke models.

**Figure 4.**
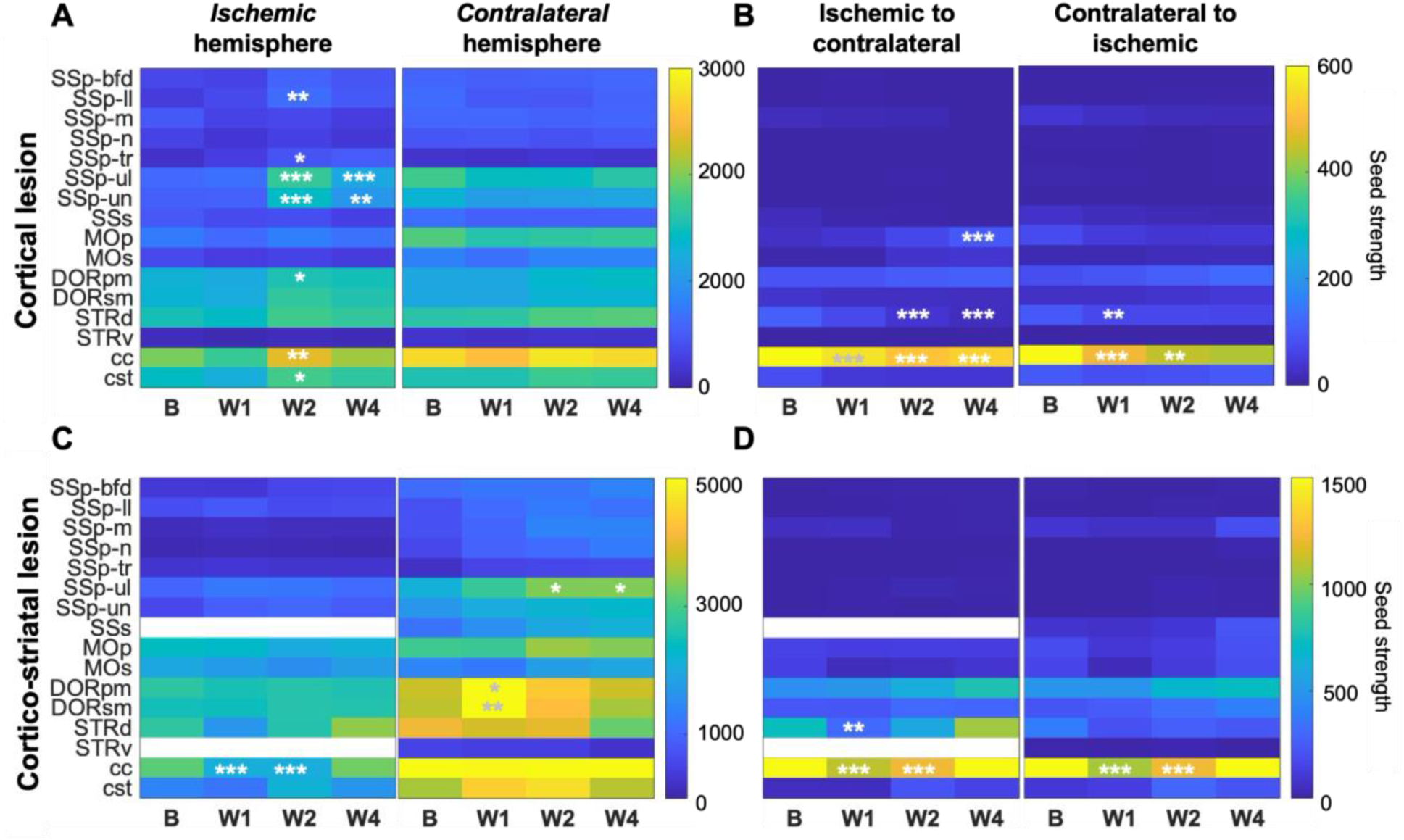
Two-dimensional representation of unilateral and bilateral seed strength changes on ischemic and contralateral hemispheres over time. Seed strength changes in selected regions of the sensorimotor network were analyzed using a mixed-effects model and post-hoc test of significant changes compared to baseline (p < 0.05 marked by single white stars, p < 0.01 with 2 stars, and p < 0.001 with 3 stars). Seed strength changes were separately analyzed for the cortical and cortico-striatal stroke model in the ischemic and contralateral hemisphere (A and C), the ischemic to contralateral and the contralateral to ischemic connections (B and D). Regions entirely affected by the stroke (∼100% affected) are marked in white and excluded from the analysis due to the absence of measurable seed strength changes.

**Cortical lesion group:** The ischemic hemisphere exhibited a significant increase in intra-hemispheric seed strengths at weeks 2 and 4 post-stroke (**Fig. 4A**). This increase was most prominent in the primary somatosensory areas: lower limb (SSp-ll, p < 0.01), trunk (SSp-tr, p < 0.05), upper limb (SSp-ul, p < 0.01), and unassigned regions (SSp-un, p < 0.001), as well as in the posterio-medial thalamus (DORpm, p < 0.05). This increase in seed strength was also linked to a significant rise in the number of fibers passing through the corpus callosum (CC, p < 0.01) and corticospinal tract (CST, p < 0.05). These changes were especially significant by week 2 after stroke. In the contralateral hemisphere, no significant changes in intra-hemispheric seed strength were observed across all regions. The strength of connections between regions of the ischemic and contralateral hemispheres showed minor changes post-stroke (**Fig. 4B**). Only by week 4 was there a significant increase in inter-hemispheric connections from the ipsilesional primary motor area (MOp) to regions on the contralateral hemisphere. However, trans-hemispheric fibers passing through the corpus callosum (cc) decreased on both the ischemic and contralateral sides, with this decline starting as early as week 1 post-stroke (p < 0.001). Almost all regions showed a decrease in connectivity, with significant reductions observed in the dorsal striatum (STRd, p < 0.01) and corpus callosum (CC) from the contralateral side to the ischemic side by week 4. However, this change was not statistically significant for the remaining regions.

**Cortico-Striatal Lesion Group:** In the ischemic hemisphere, intra-hemispheric seed strength in the sensorimotor area remained essentially unchanged after stroke (**Fig. 4C**). We observed a decrease in intra-hemispheric connections passing through the corpus callosum (CC) at weeks 1 and 2 post-stroke; however, this decrement had transitioned into an increment by week 4. In contrast, the contralateral hemisphere exhibited significant increases in intra-hemispheric seed strength. Notable increments were observed in the somatosensory-medial thalamus (DORsm, p < 0.01) and the posterio-medial thalamus (DORpm, p < 0.05) at week 1 post-stroke, while the primary somatosensory upper limb region (SSp-ul, p < 0.05) showed significant increases at week 2 and 4 post-stroke. For inter-hemispheric connections (**Fig. 4D**), seed strength from the ischemic dorsal striatum (STRd) to contralateral regions showed a significant decrease at week 1 (p < 0.01), which then shifted to an increase by week 4. Similarly, seed strength in the corpus callosum (CC) experienced a significant reduction during weeks 1 and 2 (p < 0.001), followed by an increase by week 4; however, this increase was not statistically significant compared to baseline. This trend was observed for intra-hemispheric connections passing through the corpus callosum on both the ischemic and contralateral sides.

In summary, the seed strength analysis revealed that the structural network changes were not equally present in all regions at all time points but instead restricted to specific regions and their connections in the ischemic (for cortical strokes) or contralateral hemisphere (for cortico-striatal strokes).

To quantify the ratio of regions driving the structural connectivity changes, we counted the number of regions with significant seed strength increase at specific time points (**Fig. 5**). For the **cortical lesion model**, approximately 40% of the ipsilesional regions contributed to enhanced intra-hemispheric connectivity by week 2 post-stroke, while around 20% of regions contributed to increased inter-hemispheric connectivity. Notably, in the contralateral hemisphere, about 10% of seed regions showed increased inter-hemispheric connections by week 1. However, until week 4 up to 20% of seed regions became more important.

**Figure 5.**
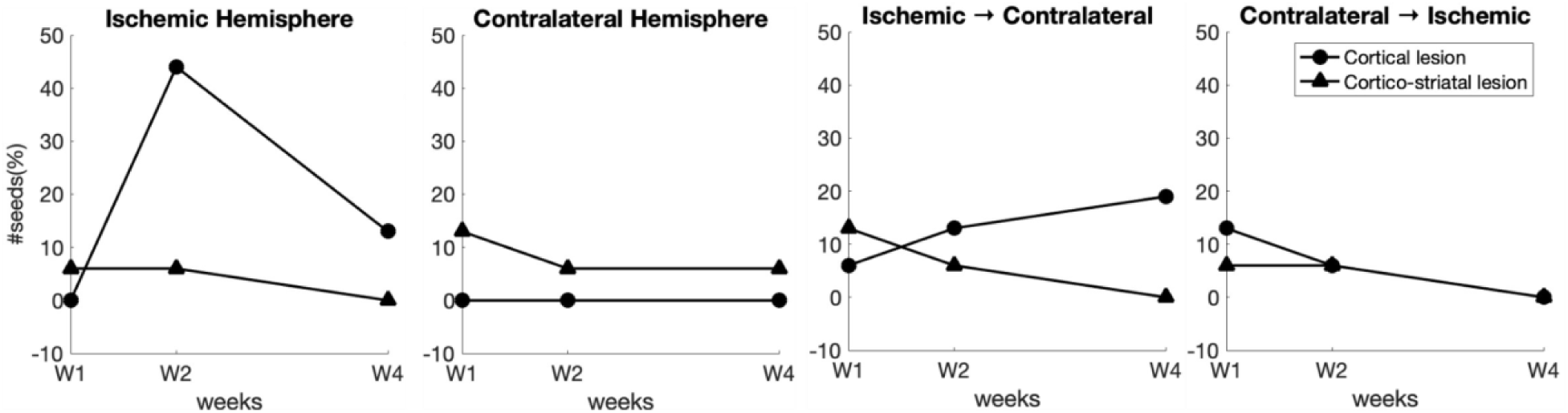
The number of regions exhibiting significantly increased seed strength within the sensorimotor network, thalamus, striatum, and selected white matter regions in the cortical and cortico-striatal stroke models.

In the **cortico-striatal lesion model** fewer regions (at all time points <10%) exhibited significant seed strength changes in the ischemic hemisphere. In the contralateral hemisphere, the percentage of regions with significant seed strength changes was at all times higher than in the cortical stroke model but still considerably smaller than the increase seen in the cortical model in the ischemic hemisphere. The inter-hemispheric connections did not substantially contribute to that increase despite week 1 with approx. 10% of regions forming significant connections from ischemic to contralateral hemisphere. However, at later times, the number of regions decreased to almost 0%, following a similar pattern in contralateral to ischemic connections.

In summary, this quantification revealed a specific pattern of seed strength changes, which were much more pronounced in the cortical lesion model in the ischemic hemisphere at week 2 and 4, mirrored by an increased importance of ischemic to contralateral connections - a dynamic not seen in the cortico-striatal model, which had more intra-hemispheric contralateral seed strength changes.

### Temporal dynamics of individual connections in somatosensory regions

In the next step of the analysis, we asked which connections forming the seed strength of a single seed actually changed over time. For that purpose, three key seed regions were selected based on their importance in behavioral improvement in experimental stroke: **SSp-ul** (upper limb somatosensory area), **SSp-ll** (lower limb somatosensory area), and **MOp** (primary motor area), and the sub-networks establishing their seed strengths were compared for each time point post stroke against baseline, separately for each hemisphere (SSp-ul in **Fig. 6**; Figure for SSp-ll, and MOp in **Online Material1^1^**). In the ischemic hemisphere, both stroke models showed a mix of increasing and decreasing connectivity changes for the SSp-ul, with only a minor set with statistically significant changes observed. In the **cortico-striatal stroke model**, connectivity between the seed region and **SSp-un** significantly increased (p < 0.05). In the **cortical lesion model**, connections to **SSp-tr** (p < 0.001), **SSp-un** (p < 0.001), and **SSp-ll** (p < 0.001) show significant increases at week 2, which largely remain significant at week 4. In the contralateral hemisphere, contrasting patterns emerged for this stroke model (**Fig. 6A**). In the **cortical lesion model**, most connections to the seed region decreased, with significant reductions in **SSp-un** (p < 0.05) at all post-stroke time points. Conversely, in the **cortico-striatal stroke model**, all connections increased relative to baseline, with significant increases in **SSp-un (**p < 0.001) and **MOp** (p < 0.05) (**Fig. 6B**).

**Figure 6.**
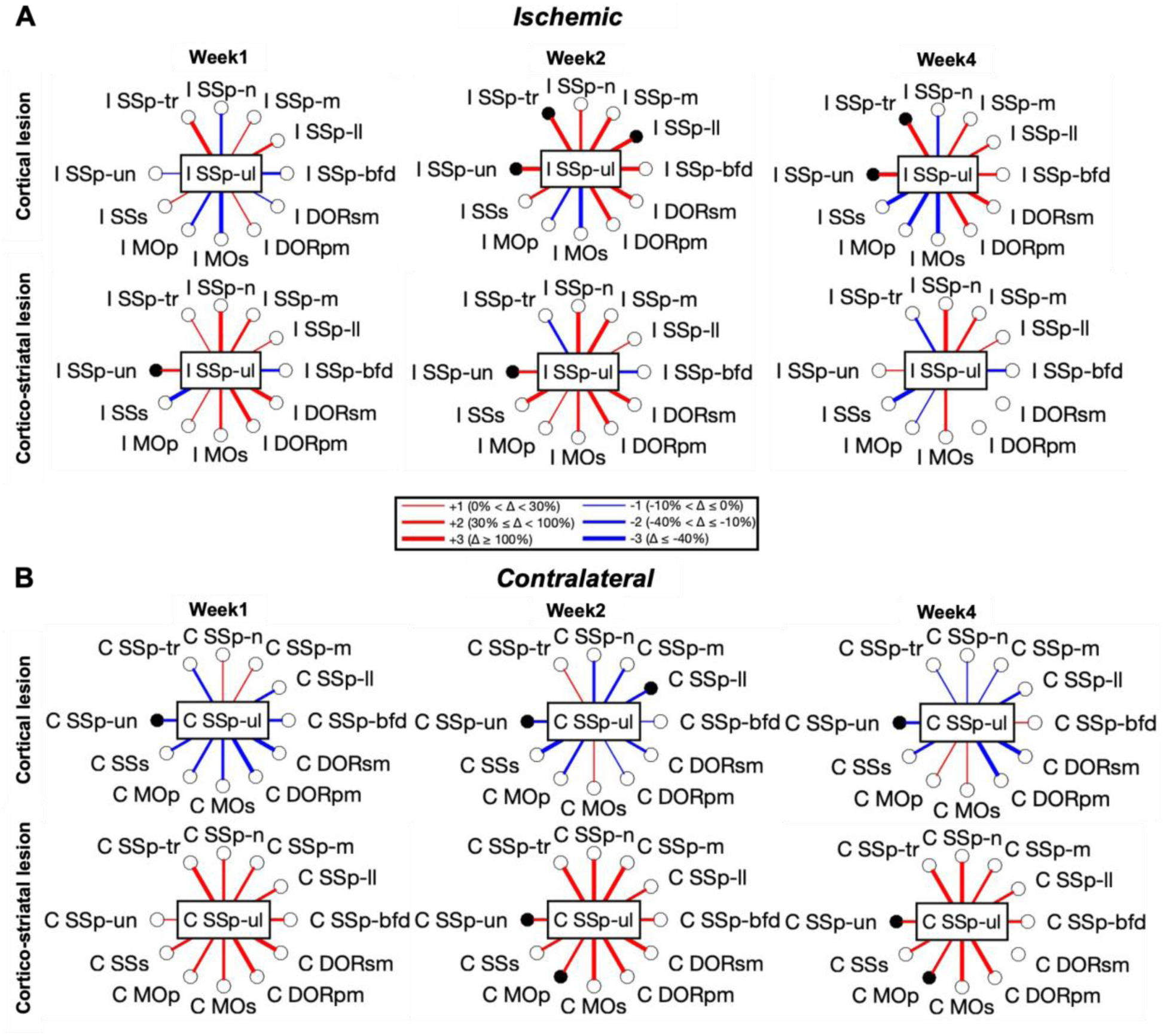
Connectivity changes in the SSp-ul network. Relative changes in fiber density compared to baseline within the ischemic (A) and contralateral hemisphere (B). The color blue indicates a weakening of connections following stroke, while red represents a strengthening of connections compared to baseline. The figure highlights the results of statistical analysis using the Linear Mixed Effects (LME) model to determine the significance of these changes (p < 0.05). Regions with significant connectivity changes are marked with solid black dots around the central node, while hollow circles represent regions with non-significant changes.

There were no statistically significant changes in inter-hemispheric connections for the upper limb somatosensory area (**Online Material^2^**). For connections from the ischemic to the contralateral side, the **cortical lesion group** exhibited a mixed pattern of changes across all time points. In contrast, the **cortico-striatal lesion group** showed a gradual increase in connectivity over time, suggesting progressive strengthening. Similarly, for connections from the contralateral side to the ischemic side, the **cortical lesion group** continues to display mixed changes, whereas the **cortico-striatal lesion group** demonstrates a general trend of increasing connectivity over time (**Online Material**^2^). In comparison to the SSp-ul, the lower limb somatosensory area (SSp-ll) and primary motor area (MOp) demonstrated distinct patterns of connectivity dynamics in both stroke models (**Online Material**^2^). In the ischemic hemisphere of the cortical lesion group, intra-hemispheric connectivity involving SSp-ll showed mixed changes, with significant increases in connections to SSp-tr and SSp-ul at week 2 and sustained increases in SSp-tr at week 4. On the contralateral side, SSp-ll connectivity predominantly decreased early post-stroke, with some connections showing recovery at later time points, including a significant increase in connectivity to the ischemic MOs at week 2 and 4. For the cortico-striatal lesion group, intra-hemispheric and inter-hemispheric SSp-ll connectivity changes generally exhibited increasing trends without significant alterations, except for a significant rise in contralateral SSp-tr connections at weeks 2 and 4.

In the cortical lesion group, MOp connectivity in the ischemic hemisphere showed significant reductions within the sensorimotor network, particularly with SSp-m. In contrast, contralateral connections exhibited a gradual increase over time, with significant increases in SSp-m and MOs at weeks 2 and 4. Conversely, connections from the contralateral MOp to ischemic sensory and thalamic regions showed progressive reductions, while connectivity to ischemic motor areas significantly increased, especially to MOs at week 2 and both MOp and MOs at week 4. For the cortico-striatal lesion group, MOp connectivity showed mixed trends, with significant reductions from the ischemic MOp to contralateral regions and predominantly significant decreases from contralateral MOp to the ischemic side, except for an increase to contralateral MOs at week 4.

These findings highlight unique temporal dynamics of individual connections for different somatosensory and motor regions and across the two lesion models, reflecting the diverse patterns of reorganization and recovery post-stroke.

To identify a pattern of structural connectivity in the two stroke models, extending beyond the single statistically significant connection changes, we determined the trends of the sub-networks forming the seed strength towards a strength increase or decrease. For this purpose, the behavior of the twelve individual connections forming the seed strength sub-network was considered. If eight or more of the contributing connections had changes of the same direction (increase or decrease), this sub-network status was considered to show a clear trend of this direction change. Otherwise, the sub-network was considered a mixed status (**Table 2**).

**Table 2.**
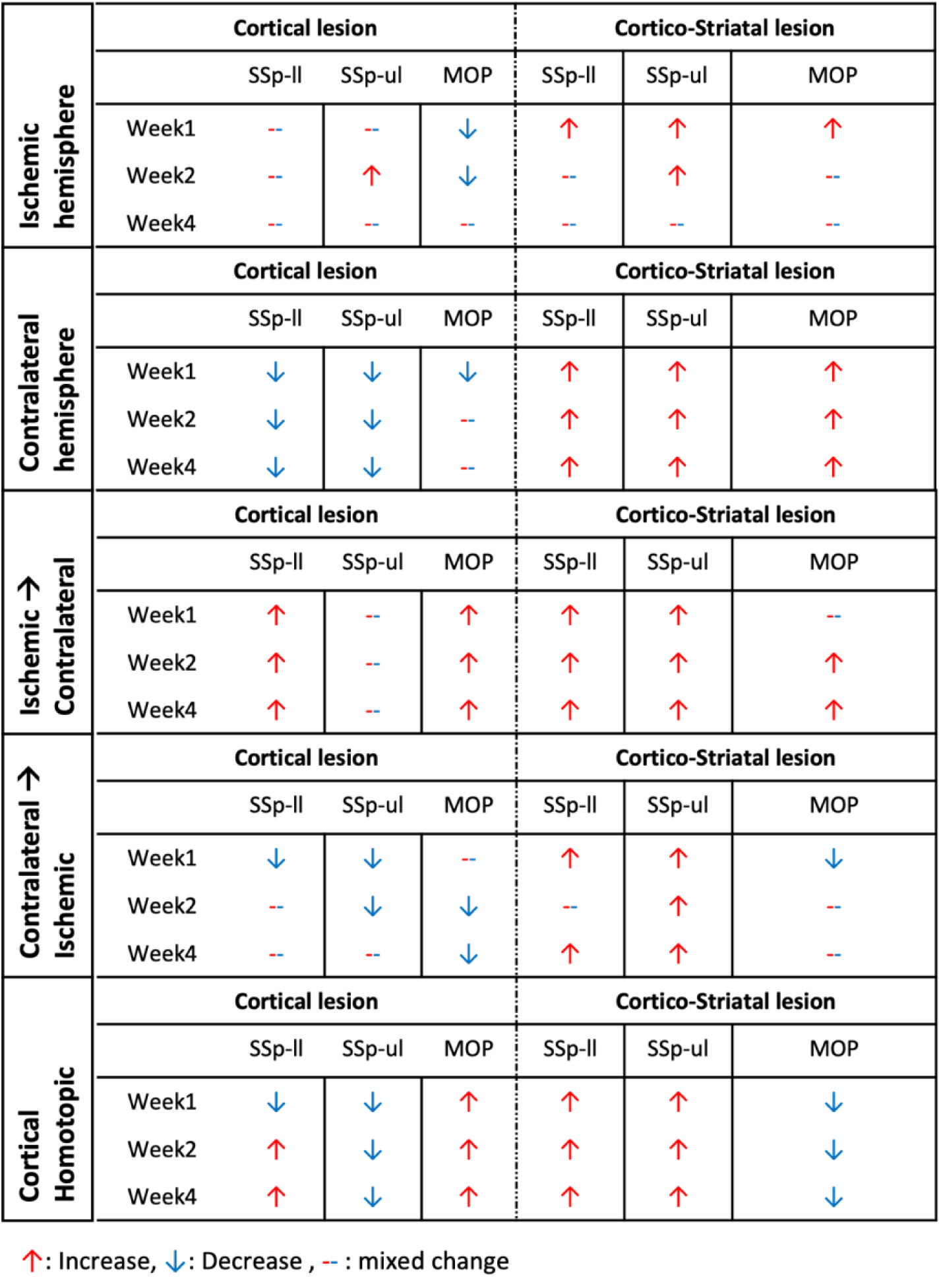
Summary of fiber density changes in key sensory-motor regions: the upper limb, lower limb, and primary motor areas.

In the **cortical lesion model**, intra-hemispheric connectivity within the ischemic hemisphere showed diverse trends, with individual connections exhibiting mixed patterns of increase and decrease compared to baseline. The motor cortex presents reduced fiber density during the first two weeks. In the contralateral hemisphere, however, most individual connections demonstrated a consistent trend of decreased connectivity relative to baseline. When considering inter-hemispheric connectivity, connections originating from seed regions in the ischemic hemisphere predominantly exhibited an increasing trend. In contrast, connections from seed regions in the contralateral hemisphere tended to show a decreasing trend compared to the baseline.

In contrast, the **cortico-striatal lesion model** displayed a more uniform pattern. The majority of individual connections, both within and between hemispheres, consistently demonstrated an increasing trend in connectivity compared to baseline, indicating an overall enhancement of structural connectivity in this model (**Table 2**).

Analyzing the individual connections, we observed distinct patterns across models. In the **cortical lesion model**, significant increases predominantly occurred in connections associated with seed regions within the ischemic hemisphere, encompassing intra- and inter-hemispheric subnetworks. Conversely, most of the significant changes originating from the contralateral hemisphere exhibited decreased fiber density. In the **cortico-striatal stroke model**, connectivity changes were largely uniform, with increased fiber density observed across most connections. This included homotopic somatosensory connections. In contrast, the homotopic motor connection showed decreased connectivity. Interestingly, the homotopic connections of the **cortical stroke model** displayed contrasting trends. Notably, the motor cortex exhibited an increase in fiber density, while the upper limb region experienced a decrease. For the lower limb region, the connectivity dynamics evolved, with initial decreases at week 1 transitioning to increases at later times (**Table 2**).

## Discussion

Diffusion MRI and to a lesser extent fiber tracking have been studied in stroke patients to correlate white matter changes related to neurodegeneration and plasticity with stroke-related motor deficits and outcomes (Boyd et al., 2017; Paul et al., 2023; Xue et al., 2024). The underlying mechanisms remain largely unknown due to a limited number of experimental stroke studies in rodents (Po et al., 2012; Anon, 2020; Aswendt et al., 2021; Sinke et al., 2021). We present to the best of our knowledge, the first longitudinal study of structural network changes after stroke in two different models, i.e., the transient MCAO and the permanent photothrombotic (PT) cortical stroke. We used deterministic fiber tracking analysis on a set of sensorimotor regions, defined by mapping the MRI data on the Allen Mouse Brain Atlas. Changes in fiber density between selected sensorimotor regions were followed over four weeks after stroke. To interpret changes in regional connectivity, seed strength, a new quantitative measure of connectivity dynamics, was introduced. This approach identified key seed regions exhibiting significant connectivity increases and highlighted differences in recovery patterns between the two stroke models. Cortical lesions demonstrated greater intra-hemispheric recovery in the ischemic hemisphere, while cortico-striatal lesions showed widespread contralateral reorganization, particularly in thalamic and sensorimotor regions. We suggest that these differences in structural reorganization mirror white matter plasticity triggered by the lesion location and topology as part of an intrinsic repair program in response to stroke. Limited white matter plasticity leading to oligodendrocyte precursor cell differentiation and myelin regeneration, as well as axonal sprouting in the peri-infarct cortex have been described previously (Carmichael et al., 2005; Cheng et al., 2024) as well as the potential of in vivo dMRI to detect subtle white matter changes, i.e., after motor-learning (Sampaio-Baptista et al., 2013) and stroke (Jiang et al., 2006).

After a transient decrease of fiber density in the ischemic hemisphere during the first week, which is repeatedly seen in similar models (van Meer et al., 2012; Jung et al., 2017), followed by a widespread ipsilateral fiber density increase and extending even to the contralateral hemisphere. Although many connectivity and seed strength changes did not reach statistical significance, different trends of the seed strength changes in the two stroke models led to the identification of key patterns of structural network dynamics over time, specific for each stroke model. A major finding in our study was the pronounced fiber density increase in various connections in both stroke models. While this appears to be in contrast with studies reporting FA decreases after stroke (Jiang et al., 2006; van Meer et al., 2012; Anon, 2020; Sun et al., 2022; Yang et al., 2022), there is ample support for substantial regenerative fiber sprouting and new post-strokeprojections (Carmichael et al., 2001; Dancause et al., 2005; Reitmeir et al., 2011). Therefore, we argue that the pronounced increase in fiber density is related to a combination of the stroke models (transient vs. permanent) and location (cortical vs. cortico-striatal). This hypothesis is supported by previous studies using transient MCAO in rats showing contralateral FA increase in the corticospinal tract (van Meer et al., 2012), ipsilateral peri-infarct FA increase (Jiang et al., 2006), increased fibers detected from a peri-infarct cortical region to corpus callosum (Jung et al., 2017), and increased fiber density in ipsilateral cortical sensorimotor areas, thalamus, striatum and white matter regions (Green et al., 2018), as well as thalamus-thalamus fiber connections (Po et al., 2012). In large but not small permanent cortical lesions (photothrombosis), we found an increase of fiber numbers between contralateral thalamus to ipsilesional cortex and homotopic cortex regions (Aswendt et al., 2021).

Increased FA and fiber tracking results have been related to GAP43, a marker of axonal sprouting (Granziera et al., 2007), and axonal orientation and myelin as measured by double Bielshowsky and Luxol fast blue stainings (Jiang et al., 2006; Jung et al., 2017). It is widely accepted that lipids, as in myelin and membranes, are a dominant source of dMRI contrast (Leuze et al., 2017), explaining changes in dMRI measures, e.g., FA. Such changes are specific for fiber density changes of white matter (Li et al., 2020) and are highly sensitive to microstructural integrity (Chang et al., 2017). However, FA is affected by the diffusion barrier of glia (Beaulieu, 2002), with a pronounced contribution in the case of gliosis after stroke (Budde et al., 2011), as well as the presence of vasogenic or cytotoxic edema, even in case of no fiber damage (Wegener et al., 2006). Opposite reports provided evidence of fiber tracking being sensitive to detect axonal reorientation also close to the glial scar (Jiang et al., 2006; Leuze et al., 2017) because of alignment between axons and reactive astrocytes (Li et al., 2005). By excluding the stroke lesion as determined at day 3 post stroke, we made sure to limit these influences on the fiber tracking. Calculating fiber densities from the resulting streamlines allows to separately monitor individual connection strengths between regions. To date, only very few studies using rodents have exploited this tract tracing approach in the context of stroke (Po et al., 2012; Jung et al., 2017; Sinke et al., 2018; Pallast et al., 2020). In the comparison of the two stroke models presented here, structural network changes of the sensorimotor cortex presented a distinctly different emphasis on degeneration and regeneration (**Fig. 7**). Cortico-striatal lesions induced a global increase of connections across hemispheres. In contrast, cortical lesions had a hemispherically distinct behavior. In the latter, connections starting from the ischemic hemisphere and extending globally showed an increased fiber density. Connections starting from the healthy hemisphere and extending globally had a decreased fiber density. The special case of cortical homotopic connections shows opposing behavior for sensory and motor regions. The behavior is the mirror-like opposite for small and large lesions. This, together with the opposite behavior of the contralateral hemisphere in the two stroke models points at a key role of the healthy hemisphere for recovery of regeneration.

**Figure 7.**
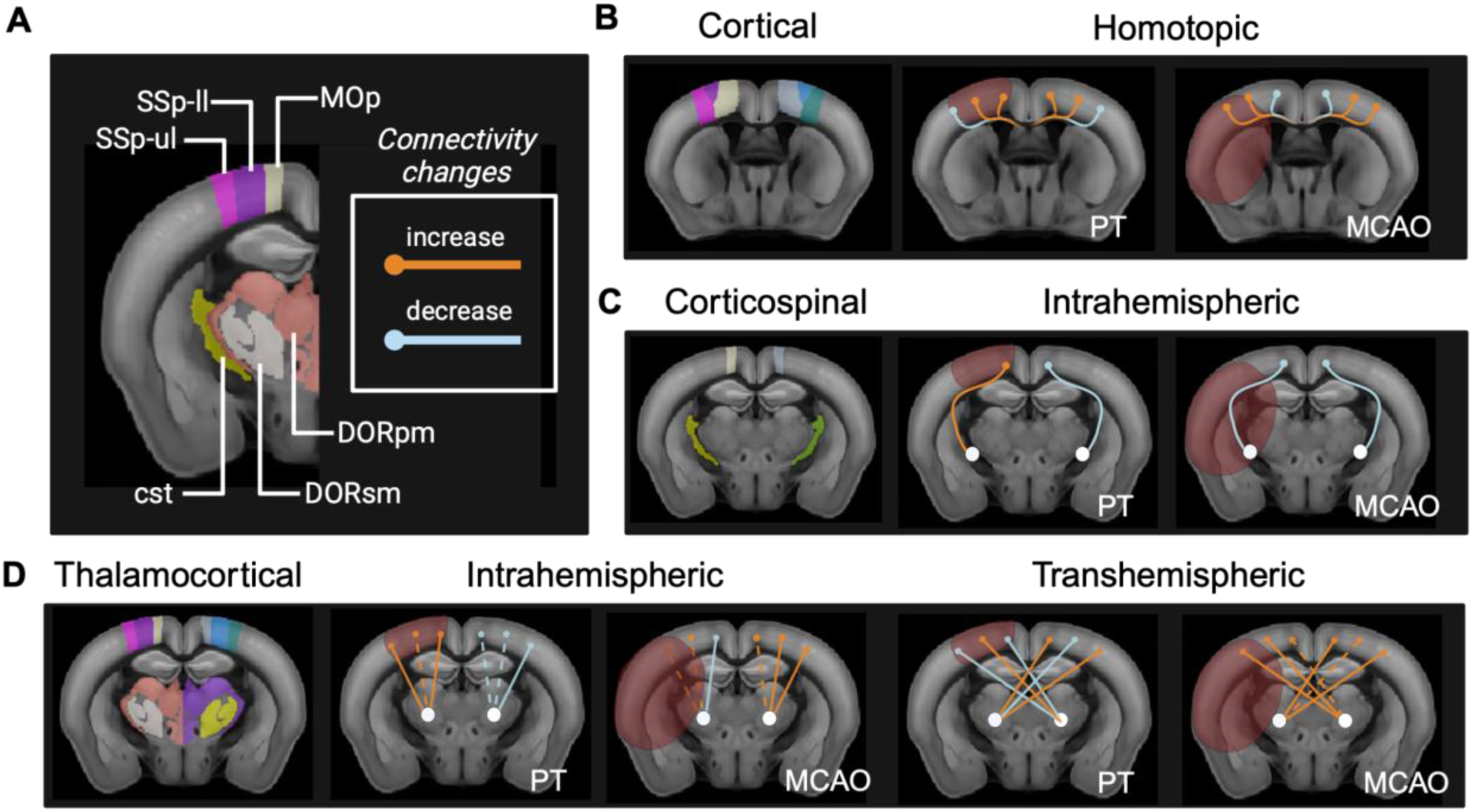
Summary of structural connectivity changes in cortical and cortico-striatal strokes. A) Illustration of the labels used for SSp-ul, SSp-ll, and MOp regions, and their connections with increased (orange) and decreased (blue) connectivity. B) Homotopic connections: In the cortical stroke model, SSp-ll and MOp demonstrated increased connectivity, whereas SSp-ul showed decreased connectivity. Conversely, in the cortico-striatal stroke model, both SSp-ul and SSp-ll exhibited increased connectivity, while MOp showed a decrease. C) Corticospinal intra-hemispheric connections: In the cortical stroke model, the ischemic hemisphere exhibited an increase in corticospinal connections, while the contralateral hemisphere showed a decrease. In contrast, in the cortico-striatal stroke model, both hemispheres demonstrated decreased corticospinal connections. D) Thalamo-cortical intra- and inter-hemispheric connections: Intra-hemispheric connections exhibited distinct patterns across the stroke models. in the cortical lesion group, the ischemic hemisphere showed partial increases in connectivity, while the contralateral hemisphere demonstrated predominantly decreases. In the cortico-striatal stroke model, most connections showed an increase in the contralateral hemisphere, whereas the ischemic hemisphere displayed a mix of increases and decreases. Transhemispheric connections also varied between the two models. Changes were mixed in the cortical lesion group and showed opposite behaviors between hemispheres. Specifically, connections from the ischemic thalamus to the contralateral MOp and SSp-ul increased, while connections to SSp-ll decreased. Conversely, connections from the contralateral thalamus to the ischemic cortex decreased to MOp and SSp-ul, but increased to SSp-ll. In the cortico-striatal stroke model, these transhemispheric connections exhibited a nearly universal increase across all regions.

The shift towards a more global fiber connection increase in cortico-striatal lesions, aligns with our previous study showing elevated fiber numbers depending on lesion size in a cortical stroke model (Aswendt et al., 2021) and a study comparing medium and large cortico-striatal strokes in rats for 14 weeks, in which large strokes showed a faster progression of FA and a significant improvement at day 70 in the ipsilesional cerebral peduncle and anterior internal capsule compared to control (van Meer et al., 2012). On the level of individual seed regions, we found no correlation between the fractional ischemic lesion territory and corresponding fiber density. However, individual connectivity changes followed a pattern: in the MOp with similar fractional lesion territory there was no significant change in both models, in SSp-ll (80% in PT, 0% in MCAO) only significant changes in PT were observed. In SSp-ul (90% in PT, 50% in MCAO) more significant changes in PT compared to MCAO were detected. The two stroke models are not only distinguished by the lesion size and topology but have distinctly different pathophysiology, which will trigger, to various degrees, aspects of spontaneous angiogenesis, neurogenesis and synaptogenesis during a recovery phase leading to different amounts of regeneration processes as reviewed in (Hossmann, 2006, 2012; Corbett et al., 2017). Photothrombosis leads to permanent vascular occlusion in the illuminated cortical territory with very little penumbra tissue produced; instead, the ischemic territory is surrounded by hyperemia (Hossmann, 2008). It is conceivable that this hyperaemia ring around the irreversibly damaged core protects the peripheral tissue and may even stimulate regeneration processes to some extent. Transient MCAO leading to cortico-striatal strokes, on the other hand, includes reperfusion and a substantial penumbra, potentially salvaged in case of successful reperfusion (Hossmann, 2012). It is conceivable that this penumbral region with viable, albeit by reduced perfusion affected tissue may act as a driver to activate regeneration processes, which have been described as a regenerative per-infarct zone of 2-mm as growth-promoting or regenerative zone for post-stroke axonal sprouting in cortical stroke in mice (Carmichael et al., 2005), which links to new patterns of intracortical projections (Carmichael et al., 2001). In concordance with our present observations where we excluded the ischemic core region from the fiber determinations, but calculated the fibers from and towards the peri-infarct tissue, long-distance projections were previously detected using axonal tracers in a cortico-striatal stroke model (Reitmeir et al., 2011).

To better understand the potential driving mechanisms, the separation of the role of the lesion volume and of the penumbra will be necessary. To elucidate the role of lesion volume alone, the permanent occlusion stroke model (Tamura et al., 1981) may be used for proximally or distally positioned occlusion of the MCA in comparison to the transient occlusion stroke model (Koizumi et al., 1986). Such variability in lesion volume, i.e., the involvement of cortex versus cortex and basal ganglia, but different pathophysiology assessed with our diffusion MRI approach and behavior, will help to further understand the clinically relevant structural connectivity changes for functional improvement.

## Data and code availability

The dataset including all raw and processed MRI data, custom scripts and code are available at https://doi.org/10.12751/g-node.ln5k5d (CC BY-NC-SA 4.0 license) according to the scheme explained in (Kalantari et al., 2023).

## Author contributions

**Fatemeh Mahani:** Methodology, Software, Formal analysis, Data curation, Writing, Visualization. **Aref Kalantari:** Methodology, Software, Data curation. **Michael Diedenhofen:** Methodology, Software, Data curation, Formal analysis. **Dirk Wiedermann:** Methodology, Software, Resources. **Gereon R. Fink:** Writing, Supervision, Funding acquisition. **Mathias Hoehn:** Conceptualization, Methodology, Formal analysis, Writing, Visualization, Supervision. **Markus Aswendt:** Conceptualization, Methodology, Resources, Data curation, Writing, Visualization, Supervision.

## Acknowledgements

This work was funded by the Deutsche Forschungsgemeinschaft (DFG, German Research Foundation): project ID 431549029–SFB 1451. The authors acknowledge technical support with in vivo experiments and MRI by Niklas Pallast, Frederique Wieters, Olivia Käsgen, Marieke Nill, Andreas Beyrau, and Ulla Uhlenküken.

## Footnotes

1 https://doi.org/10.12751/g-node.ln5k5d => Output >= Connectivity changes

2 https://doi.org/10.12751/g-node.ln5k5d => Output >= Connectivity changes

## References

Alves R, Henriques RN, Kerkelä L, Chavarrías C, Jespersen SN, Shemesh N (2022) Correlation Tensor MRI deciphers underlying kurtosis sources in stroke. Neuroimage 247:118833.

Anon (2020) Longitudinal tracing of white matter integrity on diffusion tensor imaging in the chronic cerebral ischemia and acute cerebral ischemia. Brain Research Bulletin 154:135–141.

Aswendt M, Pallast N, Wieters F, Baues M, Hoehn M, Fink GR (2021) Lesion size- and location-dependent recruitment of contralesional thalamus and motor cortex facilitates recovery after stroke in mice. Transl Stroke Res 12:87–97.

Beaulieu C (2002) The basis of anisotropic water diffusion in the nervous system - a technical review. NMR Biomed 15:435–455.

Boyd LA, Hayward KS, Ward NS, Stinear CM, Rosso C, Fisher RJ, Carter AR, Leff AP, Copland DA, Carey LM, Cohen LG, Basso DM, Maguire JM, Cramer SC (2017) Biomarkers of stroke recovery: Consensus-based core recommendations from the Stroke Recovery and Rehabilitation Roundtable. Int J Stroke 12:480–493.

Budde MD, Janes L, Gold E, Turtzo LC, Frank JA (2011) The contribution of gliosis to diffusion tensor anisotropy and tractography following traumatic brain injury: validation in the rat using Fourier analysis of stained tissue sections. Brain 134:2248–2260.

Carmichael ST (2016) The 3 Rs of Stroke Biology: Radial, Relayed, and Regenerative. Neurotherapeutics 13:348–359.

Carmichael ST, Archibeque I, Luke L, Nolan T, Momiy J, Li S (2005) Growth-associated gene expression after stroke: evidence for a growth-promoting region in peri-infarct cortex. Exp Neurol 193:291–311.

Carmichael ST, Kathirvelu B, Schweppe CA, Nie EH (2017) Molecular, cellular and functional events in axonal sprouting after stroke. Exp Neurol 287:384–394.

Carmichael ST, Wei L, Rovainen CM, Woolsey TA (2001) New patterns of intracortical projections after focal cortical stroke. Neurobiol Dis 8:910–922.

Cha J, Kim ST, Jung WB, Han YH, Im GH, Lee JH (2016) Altered white matter integrity and functional connectivity of hyperacute-stage cerebral ischemia in a rat model. Magn Reson Imaging 34:1189–1198.

Chang EH, Argyelan M, Aggarwal M, Chandon T-SS, Karlsgodt KH, Mori S, Malhotra AK (2017) The role of myelination in measures of white matter integrity: Combination of diffusion tensor imaging and two-photon microscopy of CLARITY intact brains. Neuroimage 147:253–261.

Cheng Y-J, Wang F, Feng J, Yu B, Wang B, Gao Q, Wang T-Y, Hu B, Gao X, Chen J-F, Chen Y-J, Lv S-Q, Feng H, Xiao L, Mei F (2024) Prolonged myelin deficits contribute to neuron loss and functional impairments after ischaemic stroke. Brain 147:1294–1311.

Chiu F-Y, Kuo D-P, Chen Y-C, Kao Y-C, Chung H-W, Chen C-Y (2018) Diffusion tensor-derived properties of benign oligemia, true “at risk” penumbra, and infarct core during the first three hours of stroke onset: A rat model. Korean J Radiol 19:1161–1171.

Corbett D, Carmichael ST, Murphy TH, Jones TA, Schwab ME, Jolkkonen J, Clarkson AN, Dancause N, Weiloch T, Johansen-Berg H, Nilsson M, McCullough LD, Joy MT (2017) Enhancing the alignment of the preclinical and clinical stroke recovery research pipeline: Consensus-based core recommendations from the stroke recovery and rehabilitation roundtable translational working group. Neurorehabil Neural Repair 31:699–707.

Dancause N, Barbay S, Frost SB, Plautz EJ, Chen D, Zoubina EV, Stowe AM, Nudo RJ (2005) Extensive cortical rewiring after brain injury. J Neurosci 25:10167–10179.

Dirnagl U, Iadecola C, Moskowitz MA (1999) Pathobiology of ischaemic stroke: an integrated view. Trends Neurosci 22:391–397.

Granziera C, D’Arceuil H, Zai L, Magistretti PJ, Sorensen AG, de Crespigny AJ (2007) Long-term monitoring of post-stroke plasticity after transient cerebral ischemia in mice using in vivo and ex vivo diffusion tensor MRI. Open Neuroimag J 1:10–17.

Green C, Minassian A, Vogel S, Diedenhofen M, Beyrau A, Wiedermann D, Hoehn M (2018) Sensorimotor functional and structural networks after intracerebral stem cell grafts in the ischemic mouse brain. J Neurosci 38:1648–1661.

Grefkes C, Fink GR (2014) Connectivity-based approaches in stroke and recovery of function. Lancet Neurol 13:206–216.

Hagmann P, Cammoun L, Gigandet X, Meuli R, Honey CJ, Wedeen VJ, Sporns O (2008) Mapping the structural core of human cerebral cortex. PLoS Biol 6:e159.

Hossmann K-A (2006) Pathophysiology and therapy of experimental stroke. Cell Mol Neurobiol 26:1057–1083.

Hossmann K-A (2008) Cerebral ischemia: models, methods and outcomes. Neuropharmacology 55:257–270.

Hossmann K-A (2012) The two pathophysiologies of focal brain ischemia: implications for translational stroke research. J Cereb Blood Flow Metab 32:1310–1316.

Jiang Q, Zhang ZG, Ding GL, Silver B, Zhang L, Meng H, Lu M, Pourabdillah-Nejed-D S, Wang L, Savant-Bhonsale S, Li L, Bagher-Ebadian H, Hu J, Arbab AS, Vanguri P, Ewing JR, Ledbetter KA, Chopp M (2006) MRI detects white matter reorganization after neural progenitor cell treatment of stroke. Neuroimage 32:1080–1089.

Jones TA (2017) Motor compensation and its effects on neural reorganization after stroke. Nat Rev Neurosci 18:267–280.

Jung W-B, Han YH, Chung JJ, Chae SY, Lee SH, Im GH, Cha J, Lee JH (2017) Spatiotemporal microstructural white matter changes in diffusion tensor imaging after transient focal ischemic stroke in rats. NMR Biomed 30 Available at: https://pubmed.ncbi.nlm.nih.gov/28205341/ [Accessed January 22, 2025].

Kalantari A, Szczepanik M, Heunis S, Mönch C, Hanke M, Wachtler T, Aswendt M (2023) How to establish and maintain a multimodal animal research dataset using DataLad. Sci Data 10:357.

Koizumi J-I, Yoshida Y, Nakazawa T, Ooneda G (1986) Experimental studies of ischemic brain edema. Nosotchu 8:1–8.

Leuze C, Aswendt M, Ferenczi E, Liu CW, Hsueh B, Goubran M, Tian Q, Steinberg G, Zeineh MM, Deisseroth K, McNab JA (2017) The separate effects of lipids and proteins on brain MRI contrast revealed through tissue clearing. Neuroimage 156:412–422.

Li C, Gore JC, Davatzikos C (2014) Multiplicative intrinsic component optimization (MICO) for MRI bias field estimation and tissue segmentation. Magn Reson Imaging 32:913–923.

Li M, Zhao Y, Zhan Y, Yang L, Feng X, Lu Y, Lei J, Zhao T, Wang L, Zhao H (2020) Enhanced white matter reorganization and activated brain glucose metabolism by enriched environment following ischemic stroke: Micro PET/CT and MRI study. Neuropharmacology 176:108202.

Liu J, Liu D, Pu X, Zou K, Xie T, Li Y, Yao H (2023) The secondary motor cortex-striatum circuit contributes to suppressing inappropriate responses in perceptual decision behavior. Neurosci Bull 39:1544–1560.

Li Y, Chen J, Zhang CL, Wang L, Lu D, Katakowski M, Gao Q, Shen LH, Zhang J, Lu M, Chopp M (2005) Gliosis and brain remodeling after treatment of stroke in rats with marrow stromal cells. Glia 49:407–417.

Moura LM, Luccas R, de Paiva JPQ, Amaro E Jr, Leemans A, Leite C da C, Otaduy MCG, Conforto AB (2019) Diffusion Tensor Imaging Biomarkers to Predict Motor Outcomes in Stroke: A Narrative Review. Front Neurol 10:445.

Pallast N, Diedenhofen M, Blaschke S, Wieters F, Wiedermann D, Hoehn M, Fink GR, Aswendt M (2019) Processing Pipeline for Atlas-Based Imaging Data Analysis of Structural and Functional Mouse Brain MRI (AIDAmri). Front Neuroinform 13:42.

Pallast N, Wieters F, Nill M, Fink GR, Aswendt M (2020) Graph theoretical quantification of white matter reorganization after cortical stroke in mice. Neuroimage 217:116873.

Paul T, Wiemer VM, Hensel L, Cieslak M, Tscherpel C, Grefkes C, Grafton ST, Fink GR, Volz LJ (2023) Interhemispheric Structural Connectivity Underlies Motor Recovery after Stroke. Ann Neurol 94:785–797.

Pekny M, Pekna M (2014) Astrocyte reactivity and reactive astrogliosis: costs and benefits. Physiol Rev 94:1077–1098.

Po C, Kalthoff D, Kim YB, Nelles M, Hoehn M (2012) White matter reorganization and functional response after focal cerebral ischemia in the rat. PLoS One 7:e45629.

Reitmeir R, Kilic E, Kilic U, Bacigaluppi M, ElAli A, Salani G, Pluchino S, Gassmann M, Hermann DM (2011) Post-acute delivery of erythropoietin induces stroke recovery by promoting perilesional tissue remodelling and contralesional pyramidal tract plasticity. Brain 134:84–99.

Sampaio-Baptista C, Khrapitchev AA, Foxley S, Schlagheck T, Scholz J, Jbabdi S, DeLuca GC, Miller KL, Taylor A, Thomas N, Kleim J, Sibson NR, Bannerman D, Johansen-Berg H (2013) Motor skill learning induces changes in white matter microstructure and myelination. J Neurosci 33:19499–19503.

Sinke MR, Otte WM, van Meer MP, van der Toorn A, Dijkhuizen RM (2018) Modified structural network backbone in the contralesional hemisphere chronically after stroke in rat brain. J Cereb Blood Flow Metab 38:1642–1653.

Sinke MRT, van Tilborg GAF, Meerwaldt AE, van Heijningen CL, van der Toorn A, Straathof M, Rakib F, Ali MHM, Al-Saad K, Otte WM, Dijkhuizen RM (2021) Remote Corticospinal Tract Degeneration After Cortical Stroke in Rats May Not Preclude Spontaneous Sensorimotor Recovery. Neurorehabil Neural Repair 35:1010–1019.

Sun M, Wu L, Chen G, Mo X, Shi C (2022) Hemodynamic changes and neuronal damage detected by 9.4 T MRI in rats with chronic cerebral ischemia and cognitive impairment. Brain Behav 12:e2642.

Tamura A, Graham DI, McCulloch J, Teasdale GM (1981) Focal cerebral ischaemia in the rat: 1. Description of technique and early neuropathological consequences following middle cerebral artery occlusion. J Cereb Blood Flow Metab 1:53–60.

Ueno M, Hayano Y, Nakagawa H, Yamashita T (2012) Intraspinal rewiring of the corticospinal tract requires target-derived brain-derived neurotrophic factor and compensates lost function after brain injury. Brain 135:1253–1267.

van Meer MPA, Otte WM, van der Marel K, Nijboer CH, Kavelaars A, van der Sprenkel JWB, Viergever MA, Dijkhuizen RM (2012) Extent of bilateral neuronal network reorganization and functional recovery in relation to stroke severity. J Neurosci 32:4495–4507.

van Meer MPA, van der Marel K, van der Sprenkel JWB, Dijkhuizen RM (2011) MRI of bilateral sensorimotor network activation in response to direct intracortical stimulation in rats after unilateral stroke. J Cereb Blood Flow Metab 31:1583–1587.

Wegener S, Hoehn M, Back T (2006) Ischemic Edema and Necrosis. In: Magnetic Resonance Imaging in Ischemic Stroke, pp 133–148. Berlin, Heidelberg: Springer Berlin Heidelberg.

Xue X, Wu J-J, Xing X-X, Ma J, Zhang J-P, Xiang Y-T, Zheng M-X, Hua X-Y, Xu J-G (2024) Mapping individual cortico-basal ganglia-thalamo-cortical circuits integrating structural and functional connectome: implications for upper limb motor impairment poststroke. MedComm (2020) 5:e764.

Yang L, Li M, Zhan Y, Feng X, Lu Y, Li M, Zhuang Y, Lei J, Zhao H (2022) The Impact of Ischemic Stroke on Gray and White Matter Injury Correlated With Motor and Cognitive Impairments in Permanent MCAO Rats: A Multimodal MRI-Based Study. Front Neurol 13:834329.

Yeh F-C, Verstynen TD, Wang Y, Fernández-Miranda JC, Tseng W-YI (2013) Deterministic diffusion fiber tracking improved by quantitative anisotropy. PLoS One 8:e80713.

Yushkevich PA, Piven J, Hazlett HC, Smith RG, Ho S, Gee JC, Gerig G (2006) User-guided 3D active contour segmentation of anatomical structures: significantly improved efficiency and reliability. Neuroimage 31:1116–1128.

Zanier ER, Pischiutta F, Villa P, Paladini A, Montinaro M, Micotti E, Orrù A, Cervo L, De Simoni MG (2013) Six-month ischemic mice show sensorimotor and cognitive deficits associated with brain atrophy and axonal disorganization. CNS Neurosci Ther 19:695–704.

